# Anillin tunes contractility and regulates barrier function during Rho flare-mediated tight junction remodeling

**DOI:** 10.1101/2024.11.20.624537

**Authors:** Zie Craig, Torey R. Arnold, Kelsey Walworth, Alexander Walkon, Ann L. Miller

## Abstract

To preserve barrier function, cell-cell junctions must dynamically remodel during cell shape changes. We have previously described a rapid tight junction repair pathway characterized by local, transient activation of RhoA, termed ‘Rho flares,’ which repair leaks in tight junctions via promoting local actomyosin-mediated junction remodeling. In this pathway, junction elongation is a mechanical trigger that initiates RhoA activation through an influx of intracellular calcium and recruitment of p115RhoGEF. However, mechanisms that tune the level of RhoA activation and Myosin II contractility during the process remain uncharacterized. Here, we show that the scaffolding protein Anillin localizes to Rho flares and regulates RhoA activity and actomyosin contraction at flares. Knocking down Anillin results in Rho flares with increased intensity but shorter duration. These changes in active RhoA dynamics weaken downstream F-actin and Myosin II accumulation at the site of Rho flares, resulting in decreased junction contraction. Consequently, tight junction breaks are not reinforced following Rho flares. We show that Anillin-driven RhoA regulation is necessary for successfully repairing tight junction leaks and protecting junctions from repeated barrier damage. Together, these results uncover a novel regulatory role for Anillin during tight junction repair and barrier function maintenance.

**Significance Statement:**

- Barrier function is critical for epithelial tissues. Epithelial cells maintain barrier function via tight junctions, which must be remodeled to allow for cell- and tissue-scale shape changes. How barrier function is maintained and remodeled as epithelial cells change shape remains unclear.
- The scaffolding protein Anillin is required for generating effective actomyosin contraction to reinforce damaged tight junctions; lack of reinforcement leads to repeated barrier leaks.
- These findings highlight a novel role for Anillin in tight junction remodeling and suggest that Anillin’s ability to tune the level and duration of local Rho activation affects the contractile output.

## Introduction

Epithelial morphogenesis is critical not only during development as various tissues and organs are formed, but also for the normal function and homeostasis of tissues and organs within adult organisms. Specialized cell-cell junctions between polarized epithelial cells form a selectively permeable barrier, a defining feature of epithelial tissues. The selectivity of the barrier is maintained by apical tight junctions, which occlude the space between cells via transmembrane tight junction proteins including Claudins and Occludin, which form sealing strands (Balda and Matter, 2023; Citi et al., 2024). These transmembrane tight junction proteins are linked to a junctional actomyosin belt through cytoplasmic plaque proteins such as ZO-1 (Guillot and Lecuit, 2013; Shen et al., 2011; Van Itallie et al., 2017; Varadarajan et al., 2019). Basal to tight junctions are adherens junctions, which mechanically couple cells to each other and communicate forces across the epithelium by linking to the apical actomyosin belt (Arnold et al., 2017; Guillot and Lecuit, 2013). The connection of apical cell-cell junctions to the actomyosin network is critical for maintaining barrier function, but epithelial cells must also allow for cytoskeletal and junctional remodeling during cell shape changes that regularly occur during development and homeostatic organ function (Higashi et al., 2016; Mason et al., 2016; Shindo et al., 2019; Stephenson et al., 2019). Misregulation of cytoskeletal dynamics underlies many diseases, where the resulting impairment of barrier function drives disease pathogenesis (Blaine and Dylewski, 2020; Choi et al., 2017; Ivanov et al., 2010). Despite the broad importance of epithelial cell-cell junctions, many of the potential regulatory pathways that maintain cell-cell junction integrity and barrier function during cell and tissue remodeling events remain uncharacterized.

The small GTPase RhoA is a well-known regulator of actomyosin and drives the actomyosin contractility underlying many cell shape change events during morphogenesis; RhoA also maintains the apical actomyosin belt at cell-cell junctions in stable epithelia (Breznau et al., 2015; Martin and Goldstein, 2014; Priya et al., 2015; Reyes et al., 2014). Like other GTPases, RhoA acts as a molecular ‘switch’ and is toggled between a GTP-bound active state, and a GDP-bound inactive state. Active RhoA preferentially inserts into phosphatidylinositol 4,5-bisphosphate (PIP_2_) rafts within the plasma membrane (Brill et al., 2011), and promotes downstream actomyosin contractility through actin polymerization and Myosin II activation (Arnold et al., 2017). However, Rho activation must be carefully balanced; either hyperactivation or inhibition of RhoA activity negatively affect tight junction integrity (Arnold et al., 2017; Citi et al., 2014; Quiros and Nusrat, 2014). Spatiotemporal control of RhoA activity is regulated by guanine nucleotide exchange factors (GEFs), which activate RhoA, and GTPase activating proteins (GAPs), which inactivate RhoA, as well as other effector proteins and scaffolding proteins, which can interact with RhoA and regulate its activation, deactivation, or sequestration. Specificity of functionally-distinct pathways acting downstream of RhoA is determined by the regulated expression, recruitment, and activation of these RhoA interactors at the right times and places (Arthur and Burridge, 2001; Breznau et al., 2015; Garcia-Mata and Burridge, 2007).

Our lab has characterized a mechanosensitive RhoA-dependent signaling pathway that helps tight junctions and the associated actomyosin cytoskeleton remodel to maintain junction integrity and barrier function as epithelial cells change shape (Stephenson et al., 2019; Varadarajan et al., 2019). As cell-cell junctions elongate, micron-scale breaks in the tight junction can occur. At these sites, local, transient activation of RhoA (“Rho flares”) promotes local actomyosin accumulation, which helps reinforce the tight junction proteins and repairs local leaks in the barrier (Stephenson et al., 2019). We have previously characterized several of the signals and proteins involved in this process. Junction elongation serves as a mechanical cue to initiate the Rho flare process through the activation of a mechanosensitive calcium channel, which drives intracellular calcium flashes (Varadarajan et al., 2022). p115RhoGEF localizes to sites of tight junction damage and activates RhoA, promoting localized, transient flares of RhoA activity at the sites of tight junction breaks (Chumki et al., 2022). Rho flares promote actomyosin accumulation, which locally concentrates junction proteins to repair the breaks and reinforce the junction against future damage (Stephenson et al., 2019). Rho flares are regulated with incredible spatiotemporal precision; a single Rho flare is initiated and resolved on the timescale of minutes, suggesting a role for both positive and negative regulatory elements. Indeed, activation and inactivation of RhoA is often tightly coupled in space and time *via* signaling circuits based on positive and negative feedback (Bement et al., 2024). Here, we sought to investigate additional proteins that may regulate RhoA at sites of tight junction damage and remodeling.

Anillin is a multifunctional scaffolding protein that can regulate Rho activity by interacting with Rho directly, as well as by scaffolding GEFs, GAPs, effectors, and actomyosin (Piekny and Maddox, 2010; Rezig et al., 2023). Anillin was initially characterized for its role during cytokinesis, where it concentrates at the cleavage furrow and uses its many binding domains to interact with the cytokinetic machinery including: organizing and stabilizing the actomyosin in the cytokinetic contractile ring (Field and Alberts, 1995), linking RhoA with actomyosin at the cleavage furrow (Piekny and Glotzer, 2008), anchoring the contractile ring to the plasma membrane (Straight et al., 2005), promoting asymmetric ingression of the cleavage furrow *via* its interaction with septins (Maddox et al., 2007), and recruiting p190RhoGAP-A at the end of cytokinesis to terminate RhoA activity and prevent excess contractility (Manukyan et al., 2015). It was originally thought that Anillin was only present in the cytoplasm during cytokinesis, and localizes to the nucleus during interphase (Field and Alberts, 1995). However, more recent work demonstrated that there is a pool of Anillin at cell-cell junctions and the apical membrane during interphase in *Xenopus* embryonic epithelial cells, and this pool of Anillin regulates junctional and medial-apical actomyosin (Arnold et al., 2019; Reyes et al., 2014). Moreover, other labs have identified a role for Anillin at cell-cell junctions in mammalian epithelial cells (Budnar et al., 2019; Wang et al., 2015) and in apical constriction during *Xenopus* development (Van Itallie et al., 2023). Anillin promotes junctional contractility by increasing the membrane residence time of Rho-GTP, allowing active Rho increased access to effectors (Budnar et al., 2019). Like active RhoA, Anillin preferentially interacts with PIP_2_-rich lipids (Liu et al., 2012). Furthermore, Anillin clusters PIP_2_ within the plasma membrane and transiently binds Rho, which increases the residence time of RhoA at the plasma membrane – without impeding its availability to bind to effector proteins – leading to an increase in downstream signaling output (Budnar et al., 2019; Morris et al., 2020). Given Anillin’s roles in scaffolding and regulating RhoA, actomyosin contraction, and PIP_2_ at the plasma membrane, we hypothesized that Anillin might play a role in the regulation of Rho flare-mediated tight junction remodeling.

In this study, we identify a role for Anillin in regulating Rho flare-mediated tight junction remodeling using the developing *Xenopus laevis* embryo as an intact model of the vertebrate epithelium. We use live confocal microscopy to show that Anillin localizes to Rho flares during tight junction remodeling. We demonstrate that Anillin knockdown (KD) globally reduces active Rho, and the kinetics of Rho flares are disrupted: Anillin KD Rho flares are increased in intensity but shorter in duration. Further, we find that Anillin KD reduces tight junction protein reinforcement following Rho flares. Anillin KD also reduces actomyosin contractility at Rho flares and impairs the junction shortening needed to reinforce tight junctions following damage. Finally, using a live imaging barrier assay, we show that these changes in Rho flare dynamics, local actomyosin contractility, and tight junction remodeling impair barrier function, leading to an increase in local barrier leaks. Together, this work identifies Anillin as a molecular player in the Rho flare-mediated tight junction repair pathway.

## Results

### Anillin localizes to Rho flares during tight junction remodeling

Given Anillin’s ability to interact directly with Rho and scaffold Rho regulators and actomyosin (Arnold et al., 2017; Budnar et al., 2019; Manukyan et al., 2015; Piekny and Glotzer, 2008), we first sought to investigate Anillin’s localization during Rho flares. We expressed tagged Anillin (Anillin-Halo with JF646), tagged ZO-1 (BFP-ZO1) to visualize tight junctions, and a probe for active Rho (mCherry-2xrGBD) in gastrula-stage (Nieuwkoop and Faber stage 10-11) *Xenopus laevis* embryos, and compared their localization at junctional Rho flare events using live confocal microscopy. At this stage of development, Rho flares naturally occur in the epithelium as the embryo undergoes developmentally-driven cell shape changes such as cell division that require tight junction remodeling. We found that Anillin colocalized with active RhoA at the site of Rho flares (**Figure 1A, Supplemental Video 1**). Time-lapse imaging revealed that following a decrease in ZO-1 signal, the timing of the local increase in Anillin mirrored the increase in active Rho, and ZO-1 signal was reinforced following the Rho flare (**Figure 1A**). Quantification of the intensity of active Rho and Anillin over the time-course of multiple Rho flare examples confirmed that the increase in Anillin aligned with the increase in Rho activity during Rho flares, although Anillin’s peak intensity may occur after active Rho’s peak intensity (**Figure 1, B and C, Supplemental Figure S1A**). Following the peak of active Rho and Anillin, both intensities trended back towards baseline by about 300 seconds following the start of the Rho flare (time 0) (**Figure 1C**). Together, this data demonstrates that Anillin localizes to Rho flares.

**Figure 1.**
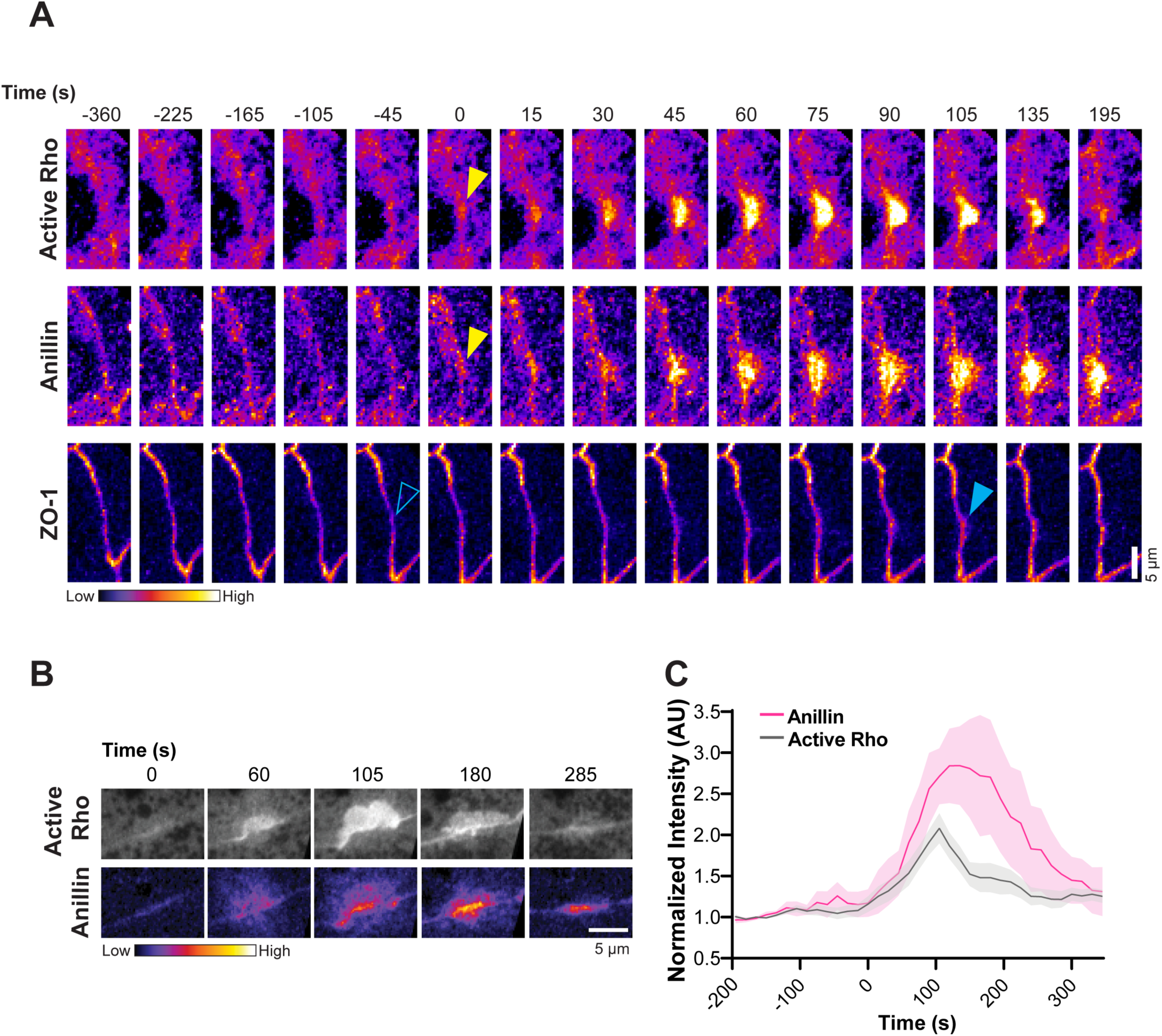
Anillin localizes to Rho flares during tight junction remodeling. (A) Time-lapse montage of Anillin (Anillin-Halo with JF 646), active RhoA probe (mCherry-2xrGBD), and ZO-1 (BFP-ZO-1), all shown in Fire LUT. Time=0 indicates the start of the Rho flare. Reduction in ZO-1 signal (open blue arrowhead) precedes Rho flare (yellow arrowhead, active Rho channel), which is accompanied by an increase in Anillin (yellow arrowhead, Anillin channel). Following the Rho flare, ZO-1 intensity is reinforced (solid blue arrowhead). (B) Anillin intensity (FIRE LUT) increases and decreases with similar timing to active Rho intensity (grayscale) over the lifetime of a Rho flare. (C) The increase in Anillin intensity (magenta line) temporally colocalizes with the increase in active Rho (gray line) during Rho flares. Quantified from (B) and additional Rho flare examples. Time 0 = the initiation of the Rho flare, shading represents SEM. n = 1 experiment, 2 embryos, 7 Rho flares.

### Anillin regulates active RhoA kinetics and tight junction reinforcement during tight junction remodeling

Because Anillin is localized at Rho flares, we next wanted to determine if Anillin plays a role in regulating active Rho and/or tight junction dynamics at sites of tight junction remodeling. We targeted Anillin for knockdown using a previously-characterized antisense morpholino oligomer (MO) targeting the 5’UTR of Anillin mRNA transcripts (Arnold et al., 2019; Reyes et al., 2014) (**Supplemental Figure S1B**) and observed the effects on Rho flares using live confocal microscopy. We observed that Anillin KD embryos exhibited decreased active RhoA signal at cell-cell junctions and the medial apical cell cortex, consistent with previous reports (**Supplemental Figure S1C**) (Arnold et al., 2019; Reyes et al., 2014). Notably, RhoA activity was more intense at Rho flare events in Anillin KD embryos relative to controls (**Figure 2, A and B, Supplemental Video 2**). To measure potential changes in active RhoA dynamics, we quantified the intensity of active RhoA over the lifetime of Rho flares in both control and Anillin KD embryos, as previously described (Chumki et al., 2022; Stephenson et al., 2019; Varadarajan et al., 2022) (**Figure 2C, Supplemental Figure S1A**). The peak intensity of active RhoA at Rho flares was significantly higher in Anillin KD embryos compared with controls (**Figure 2C’**). We calculated area under the curve for RhoA activity (from the data in **Figure 2C**). Surprisingly, we found that there was no significant difference between control and Anillin KD Rho flares, suggesting that the overall amount of RhoA activation at these sites of tight junction remodeling was similar (**Supplemental Figure S1D**). However, when we examined the change in RhoA intensity over time data, we observed that active RhoA appeared to decline faster in Anillin KD embryos compared to controls (**Figure 2C**). To quantify this, we compared the rate of decay of active RhoA signal, and found that active RhoA signal indeed decays significantly faster in Anillin KD vs. controls (**Figure 2C’’**). In summary, these results indicate that when Anillin is knocked down, Rho flare kinetics are altered such that Rho flares are more intense but shorter lived.

**Figure 2.**
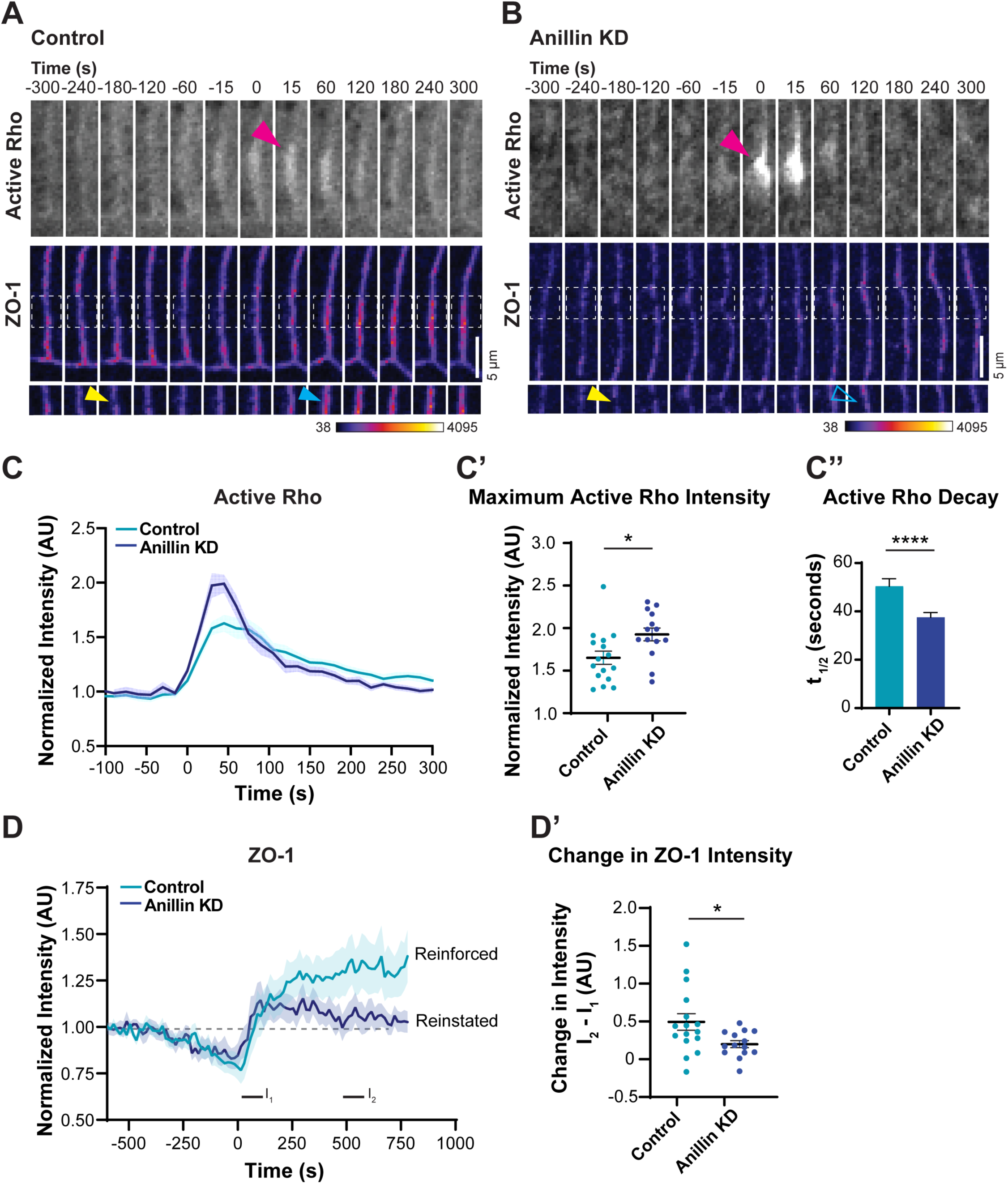
Anillin regulates active RhoA kinetics and tight junction reinforcement during tight junction remodeling. (A-B) Time-lapse montage of active RhoA probe (GFP-rGBD, grayscale) and ZO-1 (BFP-ZO-1, FIRE LUT) in control (A) vs. Anillin KD (B) embryos. White dashed boxes in ZO-1 channel indicate regions of tight junction damage and repair shown below. All images were acquired using the same microscopy settings and contrast enhanced in the same way. Anillin KD embryos show increased active Rho signal (magenta arrowhead) and ZO-1 breaks (yellow arrowhead) that fail to be reinforced following the Rho flare (open blue arrowhead). (C-D) Graphs of mean normalized intensity of active RhoA (C) and ZO-1 (D) in control (light blue lines) and Anillin KD embryos (dark blue lines) at Rho flares over time. Time 0 = the initiation of the Rho flare, shading represents SEM. n = 4 experiments, 7 embryos, 16 control flares, 14 Anillin KD flares. (C’) Scatterplot of maximum normalized intensity (averaged over 30 seconds: 15 seconds before and 15 seconds after the peak intensity for each Rho flare) of active RhoA at Rho flares is significantly higher (*p = 0.0182) in Anillin KD embryos vs. controls, quantified from (C). Error bars represent mean +/- SEM. (C’’) Active RhoA at Rho flares decays significantly faster (****p < 0.0001) in Anillin KD embryos vs. controls, quantified from (C). Bars indicate average; error bars indicate SEM. (D’) The increase in ZO-1 intensity following Rho flares is significantly smaller (*p = 0.027) in Anillin KD embryos vs. controls, quantified from (D). ZO-1 intensity is reinstated to baseline in Anillin KD, whereas it is reinforced over baseline in controls. I_1_ was measured from 0 to 30 seconds and averaged; I_2_ was measured from 495 to 555 seconds and averaged; change in intensity was calculated by subtracting I_1_ from I_2_. Error represents mean +/- SEM.

We next wondered whether these changes in active RhoA dynamics in Anillin KD embryos would affect the repair and reinforcement of tight junctions. Following Rho flares, the loss in tight junction signal is repaired, and then reinforced at the site of the break, measured by an increased intensity of tight junction proteins compared to their baseline intensity prior to the break (Chumki et al., 2022; Stephenson et al., 2019; Varadarajan et al., 2022). If Anillin-driven stabilization of RhoA activity is important for Rho flares, then we hypothesized that shorter Rho flares could have a negative effect on tight junction reinforcement. To assess tight junction reinforcement following Rho flares, we quantified ZO-1 intensity over the lifetime of Rho flares in control and Anillin KD embryos (**Figure 2, A, B, and D**). Of note, Anillin KD embryos failed to reinforce their tight junctions following Rho flares the way controls do; instead, ZO-1 intensity was reinstated to around baseline levels (**Figure 2D**). Indeed, we measured a significant reduction in ZO-1 reinforcement following tight junction breaks in Anillin KD embryos (**Figure 2D’**). We propose that the shorter, faster decaying Rho flares in Anillin KD embryos are not sufficient to promote the downstream signaling needed to reinforce tight junctions following damage. Taken together, our data demonstrate that Anillin regulates the dynamics of active Rho accumulation over time as well as effective tight junction reinforcement at sites of Rho flare-mediated tight junction remodeling.

### Anillin tunes adequate actomyosin contraction at Rho flares

During Rho flares, RhoA activation leads to the generation of actomyosin contractility to repair leaky tight junctions (Stephenson et al., 2019). Previous work had shown that Anillin KD reduces F-actin and Myosin II at cell-cell junctions and at the medial-apical cell cortex (Arnold et al., 2019; Reyes et al., 2014). Whether Anillin KD affects the actomyosin contractility generated during Rho flares remained unknown. We considered two possibilities for Anillin KD Rho flares: 1) the increase in peak active RhoA intensity could produce excess contractility that damages recovering, newly-repaired junctions or 2) the shorter duration of Anillin KD Rho flares could prevent adequate contractility from being generated to repair junctions.

To investigate changes in contractility, we first evaluated actin accumulation at the site of Rho flares using a fluorescent probe for F-actin, LifeAct-mRFP (**Figure 3A, Supplemental Video 3**). Compared to controls, Anillin KD embryos exhibited reduced F-actin at cell-cell junctions, consistent with our prior report (**Supplemental Figure S2A)** (Reyes et al., 2014). At Rho flares, the F-actin accumulation appeared confined to the junction in Anillin KD embryos, whereas F-actin was broadly accumulated around the Rho flare in controls (**Figure 3A, Supplemental Video 3**). Interestingly, line scans of F-actin intensity at Rho flares revealed that Anillin KD Rho flares have increased actin intensity over background compared with controls, mirroring the effect on active RhoA, but they show a marked reduction in the breadth of F-actin accumulation (**Figure 3C, Supplemental Figure S2B and B’**).

**Figure 3.**
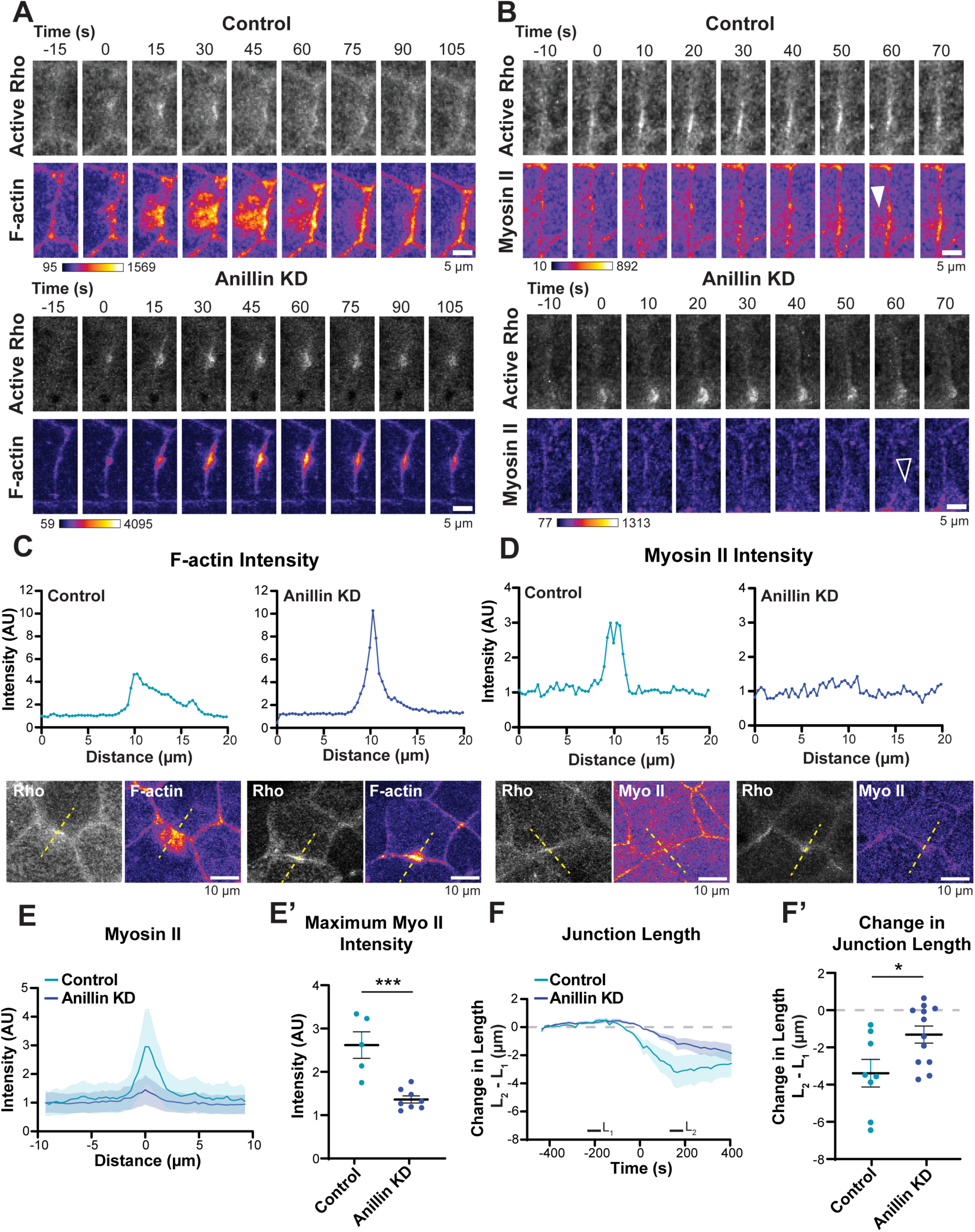
Anillin’s regulation of active Rho drives adequate actomyosin contraction at Rho flares. (A) Time-lapse montage of active Rho probe (GFP-rGBD, grayscale) and probe for F-actin (LifeAct-mRFP, FIRE LUT, adjusted as indicated on figure) in control (top) vs. Anillin KD embryos (bottom). Anillin KD embryos show an increase in local junctional actin intensity at Rho flares, but decreased breadth of actin accumulation compared with controls. (B) Time-lapse montage of active Rho probe (GFP-rGBD, grayscale) and a probe for Myosin II (SF9-mCherry, FIRE LUT, adjusted as indicated on figure), in control (top) vs. Anillin KD embryos (bottom). Control embryos show increased Myosin II at and near the junction at the site of the Rho flare (white arrowhead), white Anillin KD embryos show minimal Myosin II accumulation (open, white arrowhead) at the Rho flare. (C) Line scan analysis of F-actin intensity (line drawn through the peak of the Rho flare perpendicular to the junction, see dashed yellow lines in images below graphs). Control embryos have robust actin accumulation that spreads into the surrounding cortex vs. Anillin KD embryos, which exhibit increased actin intensity at Rho flares, but it is tightly confined at the junction. (D) Line scan analysis of Myosin II intensity (line drawn through the peak of the Rho flare perpendicular to the junction, see dashed yellow lines in images below graphs). Anillin KD embryos have decreased Myosin II accumulation at Rho flares compared to controls. (E) Averaged Myosin II intensity line scans (control, light blue line vs. Anillin KD, dark blue line), shading representing SEM. (E’) Anillin KD embryos exhibit a significant reduction in the maximum Myosin II intensity at the site of Rho flares vs. controls (***p = 0.005), quantified from (E). Error bars represent mean +/- SEM. n = 2 experiments, 3 embryos, 5 control flares, 8 Anillin KD flares. (F) Junction length measured over the time-course of Rho flares. Control embryos (light blue line) display a strong contraction that shortens junctions during Rho flares, while Anillin KD embryos (dark blue line) display less junction shortening, shading represents SEM. (F’) Change in junction length is significantly reduced in Anillin KD embryos relative to control embryos (*p = 0.0442), quantified from (F). Error bars represent mean +/- SEM. L_1_ was measured from -225 to -195 seconds and averaged, L_2_ was measured from 135 to 180 seconds and averaged, change in intensity was calculated by subtracting L_1_ from L_2_. n = 3 experiments, 6 embryos, 8 control flares, 12 Anillin KD flares.

We next evaluated Myosin II accumulation at Rho flares using a fluorescent probe for Myosin II: the SF9 intrabody (Hashimoto et al., 2015). Even more visible than the changes in F-actin, Anillin KD embryos exhibited an overall reduction in Myosin II intensity, including at cell-cell junctions (**Figure 3B, Supplemental Figure S2C, Supplemental Video 4**). Strikingly, Myosin II signal at Rho flares was barely detectable (**Figure 3D**). We quantified this effect over several examples, and Anillin KD embryos consistently showed a lack of Myosin II accumulation at Rho flares; indeed, the maximum Myosin II intensity at Rho flares was significantly reduced when Anillin was knocked down (**Figure 3E and E’**).

Next, to test the idea that the changes we observed in actomyosin accumulation and dynamics in Anillin KD embryos modulated the contractility generated at Rho flares, we measured the change in junction length over the lifetime of Rho flares (**Figure 3F**). In control embryos, junction length dramatically shortened following the Rho flare and then relaxed back toward baseline length over time (**Figure 3F**). Junction length in Anillin KD embryos did shorten to some extent following Rho flares (**Figure 3F**); however, Anillin KD junctions shortened significantly less than controls (**Figure 3F’**). Therefore, we conclude that Anillin regulation of RhoA during Rho flares is required for a robust actomyosin contractile response. These results indicate that Anillin is needed to properly tune the level of actomyosin contractility needed in order to reinforce compromised tight junctions.

### Anillin protects against barrier leaks

Because Anillin KD reduced actomyosin contractility at Rho flares and prevented the reinforcement of tight junctions (**Figures 2 and 3**), we next wanted to assay barrier function. We have previously shown that inhibiting ROCK-mediated Myosin II contractility leads to repeating Rho flares and reduced tight junction reinforcement (Stephenson et al., 2019). Additionally, targeting upstream regulators in the Rho flare tight junction repair pathway (mechanosensitive calcium channels or p115RhoGEF) impairs barrier function (Chumki et al., 2022; Varadarajan et al., 2022). To assess changes in barrier function between control and Anillin KD embryos, we used the Zinc-Based Ultrasensitive Microscopic Barrier Assay (ZnUMBA) to visualize dynamic barrier leaks, which appear as increases in FluoZin3 intensity (Higashi et al., 2023; Stephenson et al., 2019) (**Figure 4A**).

**Figure 4.**
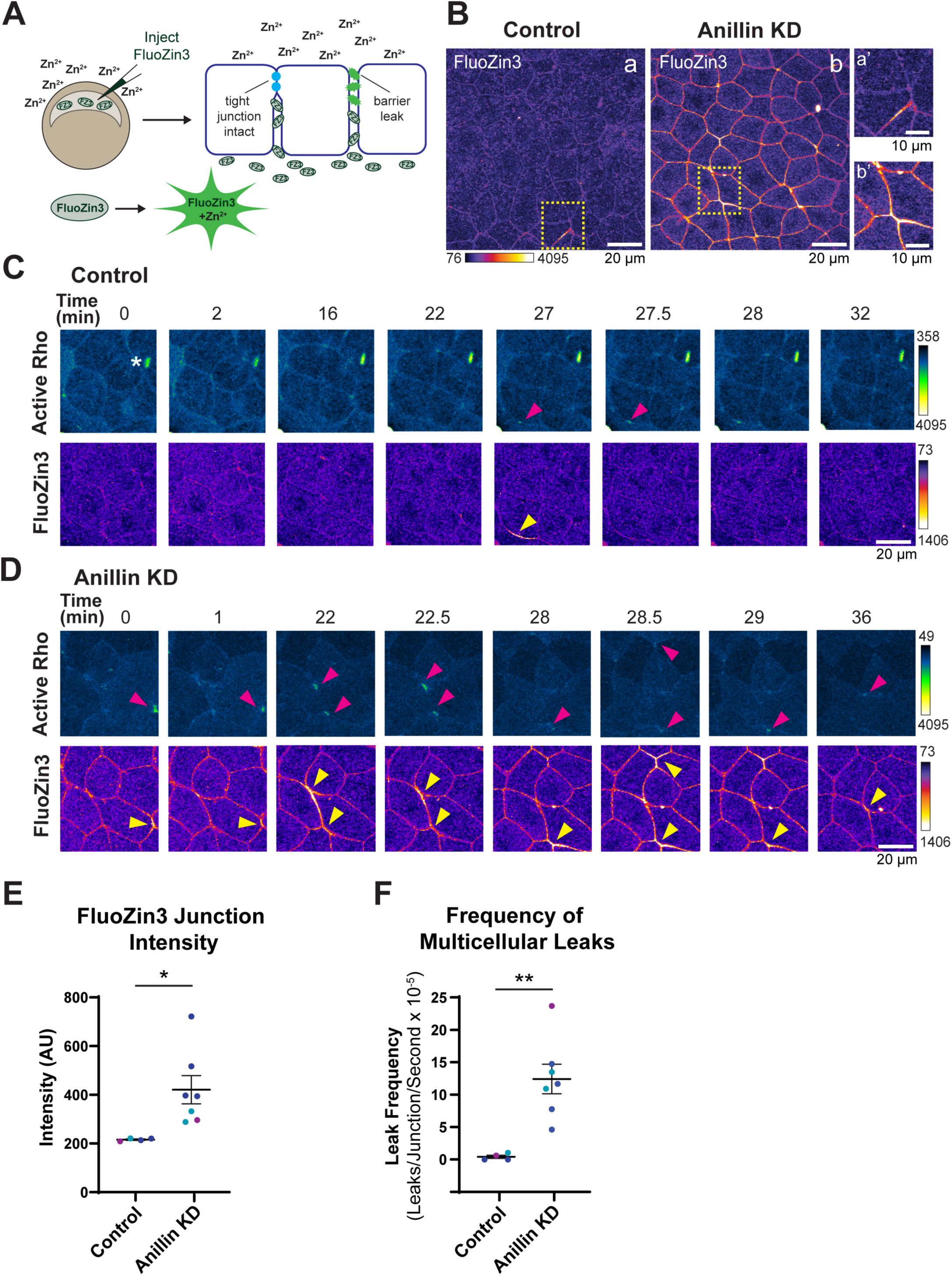
Anillin maintains barrier function and protects against repeated barrier leaks. (A) Schematic depicting Zinc-based Ultrasensitive Microscopic Barrier Assay (ZnUMBA). (B) Microscopy images showing FluoZin3 intensity in control (a) and Anillin KD embryos (b). Dashed yellow boxes indicate zoomed regions shown on the right for control (a’) and Anillin KD (b’). (C-D) Time-lapse montages of active RhoA probe (mCherry-2xrGBD, green fire blue LUT) and FluoZin3 (FIRE LUT in control (C) vs. Anillin KD embryos (D)). Pink arrowheads indicate Rho flares, and yellow arrowheads indicate barrier leaks detected by the increase in FluoZin3 intensity. White asterisk in control active Rho channel timepoint = 0 indicates bright, unchanging signal that is not relevant. Images were acquired using the same microscopy settings, and LUT levels were adjusted as listed on the figures. (E) FluoZin3 junction intensity was calculated by generating maximum intensity projections and manually tracing all junctions within the field of view from the first frame of live imaging movies and measuring their intensity. FluoZin3 junction intensity is increased in Anillin KD embryos vs controls (*p = 0.0287). Different color datapoints represent different experimental days. Error bars represent mean +/- SEM. n = 3 experiments, 4 control embryos, 7 Anillin KD embryos. (F) Frequency of barrier leaks at multicellular junctions (Multicellular Leaks/Junction/Second x 10^-5^) is increased in Anillin KD embryos vs. controls (**p = 0.0038). Different color datapoints represent different experimental days. Error bars represent mean +/- SEM. n = 3 experiments, 4 control embryos, 7 Anillin KD embryos.

In control embryos, baseline levels of FluoZin3 signal at cell-cell junctions were low, indicating that barrier function is intact (**Figure 4B**). There were occasional local barrier leaks (increased FluoZin3 signal) in controls (**Figure 4, B and C**), which triggered the Rho flare pathway, as detected by locally increased active Rho signal at the sites of FluoZin3 increases (**Figure 4C**). These leaks were quickly repaired, and FluoZin3 signal returned to baseline (**Figure 4C, Supplemental Video 5**). In contrast, Anillin KD embryos exhibited a significant increase in baseline FluoZin3 intensity at cell-cell junctions, demonstrating an overall reduction in tissue barrier function (**Figure 4, B and D, Supplemental Video 5**). Indeed, we quantified the intensity of FluoZin3 signal at junctions and found that Anillin KD embryos showed a significant increase in the FluoZin3 junction intensity (**Figure 4E**). Additionally, Anillin KD embryos exhibited a significantly higher frequency of barrier leaks happening at multicellular junctions (**Figure 4, D and F**), while there was not a statistically significant increase in local leaks along bicellular junctions (**Supplemental Figure 3B**). Taken together, we suspect that the overall decrease in barrier function in Anillin KD embryos is due to inadequate actomyosin-dependent reinforcement of compromised junctions, which predisposes tight junctions to future leaks as the tissue continues to be challenged by cell shape changes.

We noted that Anillin KD embryos often exhibit new leaks in areas that had previously been repaired or in junctions neighboring a recently-repaired junction (**Figure 4D, Supplemental Video 5**). To visualize this, we created color-coded time projections of active Rho signal from Anillin KD live imaging movies (**Supplemental Figure 3A**). These time projections revealed that in Anillin KD embryos multiple Rho flares regularly occurred along the same junction; moreover, additional flares often occurred in neighboring junctions (**Supplemental Figure 3A**). These results suggest that effective reinforcement of tight junctions following a break is essential for protecting the junction where the break occurred from further damage, as well as protecting neighboring junctions from contractility generated during the repair process. Thus, Anillin’s regulation of RhoA activation and downstream actomyosin contractility is critical for reinforcing junctions and restoring barrier function during tight junction repair.

## Discussion

Here, we have characterized a novel regulatory role for the scaffolding protein Anillin in the Rho flare tight junction repair pathway (**Figure 5**). We demonstrate that Anillin localizes to Rho flares, where it stabilizes active RhoA at the site of tight junction breaks in order to promote adequate contractility for tight junction repair (**Figures 2 and 3**). When Anillin is knocked down, we observe changes in active RhoA dynamics during Rho flares, unveiling a mechanism where Anillin tunes both the level and duration of Rho activation: in Anillin KD embryos active RhoA signal at Rho flares is more intense, but decays faster. This leads to insufficient actomyosin contractility; in particular, Myosin II does not accumulate properly at Rho flares when Anillin is knocked down. This reduced actomyosin contractile response disrupts the characteristic junction shortening normally associated with Rho flares (**Figure 3**). Without adequate contractility, damaged tight junctions are only reinstated – not reinforced – leading to an overall reduction of barrier function, as well as repeated leaks that frequently occur at multicellular junctions (**Figure 4**).

**Figure 5.**
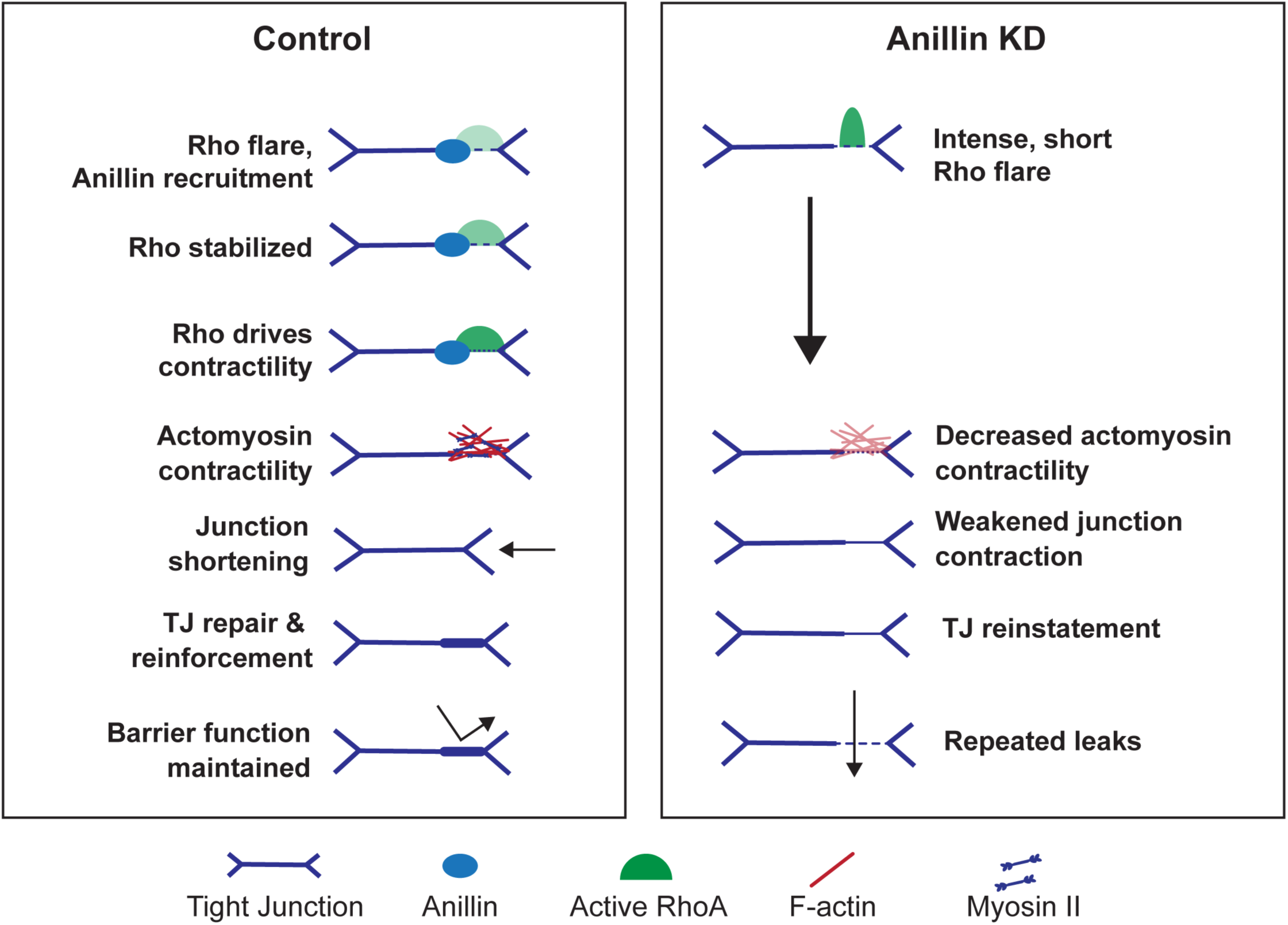
The role of Anillin in regulating actomyosin contractility at Rho flares. Schematic depicting Anillin’s role in stabilizing active RhoA at the site of Rho flares in order to promote robust downstream actomyosin contractility to repair and reinforce tight junctions, thus maintaining barrier function and protecting against additional leaks during cell shape changes. In control flares (left panel), RhoA is recruited to a break in the tight junction. Anillin is recruited to the Rho flare and stabilizes active RhoA at the site of the break to promote actomyosin contractility, which shortens the tight junction for repair and reinforcement. The reinforced junction is protected against repeating and neighboring breaks. When Anillin is knocked down, Rho flares are intense, but short, leading to insufficient actomyosin accumulation and contractility. Tight junction intensity is reinstated, but not reinforced, generating weakened areas of the junction that are prone to future breaks.

### Anillin regulates tight junction barrier function and local tight junction remodeling

Tight junction proteins form a branching strand network that regulates the selective barrier between cells, and epithelial cells must maintain and remodel this barrier even as cells change shape. Mechanisms that allow for tight junction remodeling to facilitate maintenance of barrier function in dynamic epithelial tissues are beginning to come into focus (Higashi et al., 2024; Sun et al., 2022; Varadarajan et al., 2019), with Rho flare-mediated tight junction repair being one of these.

Our lab has previously shown that Anillin regulates cell-cell junction structure. When Anillin is knocked down, tight junction proteins and actomyosin are reduced at apical cell-cell junctions (Reyes et al., 2014). In this study, we confirm that Anillin KD decreases ZO-1 intensity at cell-cell junctions, and we present new data showing that ZO-1 signal is not reinforced following Rho flares in Anillin KD embryos (**Figure 2**) as well as new functional data showing that loss of Anillin affects barrier function (**Figure 4**). One possible explanation for the lack of tight junction reinforcement is a decrease in available transmembrane tight junction proteins (e.g., claudins or Occludin) or cytoplasmic plaque proteins (e.g., ZO-1) associated with cell-cell junctions or in the nearby cytoplasm. In support of this, we see that the baseline junctional intensity of FluoZin3 is increased in Anillin KD embryos, and the spread of FluoZin3 signal often extends across multiple junctions at multicellular leak sites, suggesting that tight junctions are weaker overall when Anillin is knocked down (**Figure 4**). We speculate that this result could indicate a change in the structure of the tight junction strand network due to Anillin not properly scaffolding Rho and actomyosin at cell-cell junctions. For example, Anillin KD could decrease the number or branching pattern of tight junction strands, thus impacting baseline barrier function (Claude and Goodenough, 1973; Saito et al., 2021). Additionally, reduced junctional F-actin in Anillin KD embryos could impact the local concentration of ZO-1 near tight junctions (Shen and Turner, 2005). Further, ZO-1 has been shown to drive tight junction formation via phase separation, which is concentration dependent (Beutel et al., 2019; Schwayer et al., 2019).

Anillin KD also decreases junctional active RhoA, which is important for orchestrating proper tight junction structure and function through regulating the accumulation and contractility of actomyosin at junctions. Recent work using ZnUMBA demonstrated that the tight junction strand network is only weakly reinstated at the beginning of the Rho flare, creating a temporary seal, and that the strand network is further strengthened after actomyosin contraction has completed (Higashi et al., 2023). Due to the importance of actomyosin contractility in maintaining, repairing, and reinforcing tight junctions, we speculate that the defects in barrier function we observe in Anillin KD embryos is the cumulative effect of ineffective tight junction reinforcement from weakened actomyosin contractility, versus an overall change in tight junction strand structure. However, the observed effect could be due to a combination of several factors. Future studies with superresolution microscopy or freeze fracture electron microscopy will be needed to gain strand-level resolution.

### Recruitment of Anillin to Rho flares

An open question is how Anillin is recruited to Rho flares. When we measured the intensity of Anillin and active RhoA over the lifetime of Rho flares, we observed that both increase in intensity with similar timing, with the peak Anillin intensity appearing to be shortly after the peak of active RhoA (**Figure 1**). This timing supports the idea that Anillin may be recruited to Rho flares through interactions with Rho or other proteins in the Rho flare pathway. One possibility is that Anillin is recruited by active RhoA, similarly to how Anillin and active RhoA reciprocally stabilize each other during cytokinesis (Piekny and Glotzer, 2008; Piekny and Maddox, 2010). Anillin could also be recruited primarily through its direct interactions with F-actin and Myosin II. Given Anillin’s known interactions with RhoGEFs and RhoGAPs during cytokinesis (Frenette et al., 2012; Manukyan et al., 2015), such interactions could also potentially recruit Anillin to Rho flares. Additionally, as a multivalent scaffold protein, it is possible that Anillin participates in phase separation at cell-cell junctions; indeed, the Anillin-related protein Mid1p in fission yeast includes an intrinsically disordered region that undergoes phase separation (Chatterjee and Pollard, 2019). Future studies using mutants of Anillin that disrupt interactions with various binding partners will be needed to investigate these possibilities.

Anillin is also known to interact with phospholipids in the plasma membrane through its C2 and PH domains, with a higher affinity for negatively charged phospholipids like PIP_2_ (Liu et al., 2012; Sun et al., 2015). During cytokinesis, PIP_2_ is enriched in the cleavage furrow, and Anillin is targeted to the cleavage furrow by interactions with PIP_2_ (Liu et al., 2012). Similarly, at cell-cell junctions, Anillin is able to concentrate PIP_2_ rafts in the plasma membrane in order to extend RhoA’s residence time at the plasma membrane, increasing the opportunity for Rho to interact with effector proteins to generate more contractility (Budnar et al., 2019). Future studies should investigate whether Anillin’s interaction with PIP_2_ are needed for Anillin’s recruitment to Rho flares and/or whether Anillin facilitates concentration of PIP_2_ at Rho flares. PIP_2_ rafts could stabilize active Rho at Rho flares and also generate docking sites for Rho regulators. For example, many GEFs and GAPs contain phospholipid binding regions, and could be recruited or stabilized by PIP_2_-rich lipid rafts (Heraud et al., 2019). Moreover, PIP_2_ interactions can affect the Rho GTPase substrate preferences of RhoGAPs (Ligeti et al., 2004; Mandal, 2020). Additional studies are needed to better understand how regulation of phospholipids contributes to tight junction remodeling. In particular, future work should evaluate the localization and potential accumulation of PIP_2_ during Rho flares as a potential mechanism for protein recruitment and stabilization.

### ‘Goldilocks’ contractility at Rho flares

Although we have characterized several mechanisms regulating the local, transient activation of Rho needed for Rho flare-mediated tight junction remodeling, a number of open questions remain. Here, we have revealed new insights about Anillin’s role in tuning the intensity and duration of Rho activation in order to elicit an actomyosin response that can reinforce damaged tight junctions. However, Rho flares are spatially localized and temporally short lived, suggesting that negative regulation of RhoA is critical for successful tight junction repair. Contractility at Rho flares needs to be tightly regulated for several reasons: 1) to keep the RhoA activation spatiotemporally confined, 2) to generate adequate contractility for tight junction repair, and 3) to terminate RhoA activity after adequate contractility has been reached.

#### Spatiotemporal confinement

RhoGAPs inactivate RhoA by stimulating its intrinsic GTPase activity and are critical regulators of the spatiotemporal patterning of RhoA activity (Bement et al., 2024). RhoGAPs play an important role in the confinement of RhoA signal, and in order to do so, RhoGAPs are often coupled or complexed with RhoGEFs (Bement et al., 2024; Muller et al., 2020). The early recruitment of Anillin to Rho flares (**Figure 1**), along with our finding that Anillin KD Rho flares exhibit an increase in peak active RhoA intensity (**Figure 2**), suggests that Anillin could be scaffolding a RhoGAP in order to spatiotemporally confine the amount of active RhoA at the site of tight junction breaks. Of note, Anillin’s binding partner during cytokinesis, p190RhoGAP-A, localizes to Rho flares (**Supplemental Figure 2, A-C**), making it an excellent candidate for future investigation. Interestingly, p190RhoGAP-A appears to increase in intensity at the same time as Anillin and active RhoA (**Supplemental Figure 2, B and C**). We hypothesize that this early recruitment of a RhoGAP could potentially confine the spatial distribution and level of active RhoA at Rho flares.

#### Adequate contractility

In addition to its important role during cytokinesis, Anillin also regulates both junctional and medial-apical actomyosin (Arnold et al., 2019; Reyes et al., 2014). Our current study reveals that in addition to overall reduction of actomyosin at cell-cell junctions, the organization of both F-actin and Myosin II are disrupted at Rho flares when Anillin is knocked down (**Figure 3**). This disruption leads to insufficient contractility and inadequate tight junction repair, leading to barrier function defects (**Figures 3 and 4**). Indeed, previous work has shown that an intermediate expression level of crosslinkers like Anillin is critical for efficient contraction of the cytokinetic contractile ring: either too much or too little impairs contraction (Descovich et al., 2018). Interestingly, we noted differences in the effects of Anillin KD between F-actin and Myosin II (**Figure 3**). Actin patterning changes more closely aligned with the changes in active RhoA observed in Anillin KD embryos. Surprisingly, Myosin II intensity at Rho flares was nearly abolished when Anillin was knocked down. These results suggest that length of time RhoA is active may a critical driver of a robust contractile response, and longer RhoA activation is needed to activate Myosin II accumulation. In addition, it is known that Anillin binds to Myosin II and spatially regulates its contractility during cytokinesis (Piekny and Glotzer, 2008; Straight et al., 2005), suggesting that Anillin may also be needed to bind and stabilize Myosin II at Rho flares. Taken together, we speculate that Myosin II may be more dependent on either Anillin scaffolding at Rho flares or the length of time that active RhoA is stabilized at Rho flares, although more research is needed to clarify these possibilities. Based on our data, it seems then that the total amount of Rho activity during Rho flares is not the defining factor for adequate contractility when re-establishing tight junctions, but rather sustained Rho activity over time is needed to ensure the robust accumulation of F-actin and Myosin II contractility (**Figures 2 and 3**).

#### Termination of contractility

At Rho flares, RhoA activity rapidly increases locally – but transiently – as Rho activity goes back to baseline within hundreds of seconds. This leads us to surmise that there must be a mechanism to terminate Rho activity once adequate contractility to repair the tight junction has been reached. Excess Rho-mediated contractility has a negative effect on cell-cell junctions (Arnold et al., 2017; Citi et al., 2014; Quiros and Nusrat, 2014). Indeed, too much contractility during Rho flares could negatively impact tight junction repair and barrier function by placing mechanical strain on an already weakened junction. Thus, a ‘goldilocks’ amount of contractility is needed: enough to adequately repair the tight junction strand network without further damaging weakened tight junctions. How do cells know that adequate contractility has been reached? In the literature, Anillin has been shown to participate in both positive and negative regulation of RhoA (Budnar et al., 2019; Manukyan et al., 2015). Our data suggests that Anillin may coordinate positive and negative regulation of RhoA during Rho flares: Anillin KD leads to an increase in Rho flare intensity (suggesting Anillin is needed for negative regulation) while also reducing the duration of Rho flares (suggesting Anillin is needed for positive regulation) (**Figure 2**). One explanation could be that Anillin acts as a scaffold during Rho flares by clustering PIP_2_ to locally increase RhoA activity at the membrane, recruiting RhoGEFs and/or RhoGAPs, and enhancing local F-actin and Myosin II accumulation. Formation of signaling circuits that couple positive and negative feedback is a motif cells frequently used to pattern Rho activity (Bement et al., 2024). Another possibility is that once a certain level of contractility has been reached, Anillin mechanosensitively interacts with a RhoGAP, which then negatively regulates RhoA. In support of this idea, Anillin interacts with p190RhoGAP-A in a tension-dependent manner at the end of cytokinesis to terminate RhoA activity, and targeting contractility with blebbistatin abolishes this interaction (Manukyan et al., 2015).

In this study, we identified a novel role for Anillin in the Rho flare-mediated tight junction repair pathway where Anillin stabilizes active RhoA in order to promote sufficient actomyosin contractility for the repair and reinforcement of tight junctions and maintenance of barrier function. Our work reveals the importance of tuning the level and duration of local actomyosin contractility for effective barrier maintenance.

## Materials and Methods

### *Xenopus laevis* embryos and microinjections

All experiments conducted using *Xenopus laevis* embryos were in compliance with the University of Michigan Institutional Animal Care and Use Committee and the U.S. Department of Health and Human Services Guide for the Care and Use of Laboratory Animals. The day prior to egg collection, female *Xenopus laevis* frogs (Xenopus 1) were induced to hyper-ovulate using human chorionic gonadotrophin (HCG; MP Biomedicals, #198591). The following day, eggs were collected from female frogs and fertilized *in vitro*. Fertilized embryos were de-jellied approximately 30 minutes after fertilization and transferred to 0.1X MMR (10 mM NaCl, 0.2 mM KCl, 0.2 mM CaCl_2_, 0.1 mM MgSO_4_, and 0.5 mM HEPES, pH ≥ 7.4). Once embryos developed to the 2- or 4-cell stage, they were microinjected with the desired mRNAs and/or morpholinos. Cells were injected once per 4 cells or twice per 2 cells. For Anillin, ZO-1, and active Rho colocalization data, each 5 nL of injection volume contained: 70 pg of Anillin-Halo and 5 µM Janelia Fluor® 646 HaloTag® Ligand (Promega, #GA1120), or 70 pg of Anillin-3xGFP, 70 pg of BFP-ZO-1, and 45 pg of mCherry-2xrGBD. For Anillin KD Rho flare and ZnUMBA imaging, albino *Xenopus laevis* embryos were used. Embryos were injected with either Anillin morpholino (needle concentration of 1.25 mM), or a volume of water equal to the volume of Anillin morpholino for controls, and both were co-injected with mRNA constructs: 70 pg of BFP-ZO-1, 80 pg of GFP-rGBD or 45 pg mCherry-2xrGBD, and 45 pg of LifeAct-mRFP or mCherry-SF9, per 5 nL injection volume. Injected embryos were then incubated in 0.1X MMR and allowed to develop for approximately 16-20 hours at 15°C until they reached gastrula stage (Nieuwkoop and Faber stages 10–11). For ZnUMBA experiments, gastrula-stage embryos were injected once into the blastocoel with 10 uL with a concentration of 1mM FluoZin3 (Invitrogen, #F24195) the day after primary injections and were allowed to rest for at least 10 minutes prior to live imaging.

### DNA constructs

Anillin-mNeon was cloned by PCR amplifying Anillin from pCS2+/Anillin-3xGFP plasmid (Arnold et al. 2019) and inserting it into a digested pCS2+/C-terminal mNeon vector (EcoR1 and Xba1) using Gibson cloning (catalog #). PCS2+/mNeon-p190RhoGAP-A was generated by amplifying p190RhoGAP-A.L (NP_001084674) from PCS2+/p190 using PCR and cloned into a digested N-terminal mNeon PCS2+ vector (BamH1 and Xho1) using Gibson cloning. Constructs were initially verified by sanger sequencing (GENEWIZ) to verify short regions of interest, and then sent to Plasmidsaurus for full-plasmid sequencing. All other DNA constructs were previously described constructs utilized from the Miller lab strain collection: pCS2+/BFP-ZO1 (Stephenson et al., 2019), pCS2+/mCherry-2xrGBD (Davenport et al., 2016), pCS2+/LifeAct-mRFP (Higashi et al., 2016), and pCS2+/mCherry-SF9 (Landino et al., 2023).

### Anillin morpholino

A custom morpholino oligomer (Morpholino sequence: 5’ TGGCTAGTAACTCGATCCTCAGACT 3’) that targets the 5’UTR of Anillin mRNA transcripts was ordered from GeneTools as previously described (Arnold et al., 2019; Reyes et al., 2014).

### mRNA preparation

DNA was linearized using either Not1 or Kpn1 (in the case of ZO-1 and Anillin constructs). Linearized DNA was then transcribed using the mMessage mMachine SP6 Transcription Kit (Fisher, #AM1340). Transcribed mRNA was purified using the RNeasy Mini Kit (Qiagen, #74104) and stored at -80°C until use.

### Live imaging

Live imaging videos of gastrula-stage *Xenopus* embryos was acquired using an Olympus FluoView 1000, and 60X super-corrected Plan Apo N 60X OSC objective (NA = 1.4, working distance = 0.12 mm), with mFV10-ASW software. Embryos were mounted as previously described (Reyes et al., 2014). For control and Anillin KD Rho flares, including F-actin and Myosin II imaging, embryos were imaged with a scanning speed of 2 µs/pixel, a 512×312 field of view, 1.5X zoom, capturing 6 or 8 of the most apical slices of the animal cap of the embryo, with a step size of 0.5 µm, and time interval of 15 seconds. Colocalization of BFP-ZO-1, mNeon-p190RhoGAP-A, active Rho (mCherry-2xrGBD), Anillin-Halo (JF 646) and were imaged with a scanning speed of 2 µs/pixel, a 512×312 field of view, 1.5 zoom, capturing 3 or 4 of the most apical slices of the embryo, with a step size of 0.5 µm. For ZnUMBA imaging, embryos were mounted in 0.1X MMR containing 1 or 2 mM ZnC_l2_, as previously described (Stephenson et al., 2019).

### Image analysis

#### Rho flare quantification

Rho flares were quantified as previously described (Chumki et al., 2022; Stephenson et al., 2019; Varadarajan et al., 2022) with the exception that the circular ROI measured 0.99 µm. Rho flares were identified as a local active RhoA increase for at least 2 out of 4 frames with a 5% increase from initial baseline signal. Rho flare intensity was normalized by dividing the flare intensity values by the intensity at a reference junction not experiencing a flare. Each flare and reference junction were measured in triplicate and then averaged. Anillin, F-actin, and Myosin II at Rho flares were quantified using the same method as Rho flares, with a larger circular ROI measuring 4.05 µm. Intensity was then plotted over time with SEM shown as shaded regions above and below the average line.

#### Area under the curve

Area under the curve (AUC) was generated using Graph Pad Prism 10.2 with the following parameters: Baseline set to 1.0, and peaks that were under 10% were excluded from analysis. Individual AUC were quantified from the intensity plots of individual Rho flares (active Rho or F-actin signal) for control and Anillin KD.

#### Maximum intensity of Rho flares

Peak intensity was quantified for active RhoA by averaging the intensity values measured from 60s to 105s for each Rho flare quantified.

#### Active Rho decay

Decay was calculated by aligning Rho flares by the maximum active RhoA intensity reached as the starting value, normalized to 1 by dividing all values by the baseline intensity value at t=0, and using GraphPad Prism to calculate exponential decay with Y0=1 as a baseline.

#### Change in ZO-1 intensity

Change in ZO-1 intensity was calculated by averaging the intensity values from 0-30 seconds (I1) and averaging the intensity values from 495-555 seconds (I2) and then subtracting I1 from I2.

#### Line scan quantification of F-actin and Myosin II intensities at Rho flares

A 5 pixel-wide line measuring 20 μm was placed through the center of the Rho flare, perpendicular to the junction, using the ZO-1 signal as the center point of the junction and the highest intensity frame of the Rho flare. Line scan values were aligned via the highest value sustained for three frames, averaged, and normalized to begin at 1 by dividing individual values by the average of first 5 µm as a baseline. Only bicellular Rho flares were quantified. Rho flares were excluded if they were partially out of the field of view, repeating, the line passed through other junctions, or connected to a cleavage furrow.

#### Maximum intensity of Myosin II

Maximum intensity was calculated by averaging the highest intensity and two surrounding increments (a total of 0.993 µm) calculated from line scan measurements.

#### Change in junction length

Junction length was measured using a segmented 3 pixel-wide line to trace the junction from vertex to vertex starting 435 seconds before the flare and measuring the junction every 10 frames until 465 seconds after the Rho flare (time 0 = peak of RhoA activity during the Rho flare). These measurements were added to the ROI manager in ImageJ and the plugin Time Lapse, then LOI interpolator was used to generate ROIs for all of the frames between. Each junction was measured in triplicate and averaged. Change in length was calculated by subtracting the average length of 135 to 180 seconds (L_2_) from the average of -225 to -195 seconds (L_1_).

#### ZnUMBA: leak frequency quantification

Barrier leaks were counted by areas of FluoZin3 intensity that increased over baseline intensity revealed by a change in color using the Fire LUT applied in ImageJ. This count was then normalized by the number of junctions present in the field of view and the duration of the movie (in seconds).

#### ZnUMBA: FluoZin3 junction intensity analysis

The first frame of each video was used to create a maximum intensity projection (8 z slices). Cell-cell junctions were manually traced using a circular ROI that was 3 pixels wide using the ImageJ brush tool and measured to determine the total intensity of FluoZin3 signal at junctions. All videos were acquired using the same laser power for the FluoZin3 channel.

#### ZnUMBA: Temporal color-code projection

The temporal color code projection was made by pseudo-coloring the active RhoA channel every ten frames, and projecting these frames on top of one another to create a single image where each ten-frame time projection is represented by a different color. Rho flares that occur in the same location through different time points will overlap multiple colors, producing an additive color result (i.e., red and green overlapped to produce yellow).

#### Statistical analysis

All quantifications of control and Anillin KD embryos were compared using an unpaired Student’s T-test, calculated through Graph Pad Prism 10.2.

## Abbreviations

GAP: GTPase-activating protein;
GBD: GTPase-binding domain;
GEF: guanine nucleotide exchange factor;
KD: knockdown;
LUT: Lookup Table;
MMR: Marc’s Modified Ringers;
MO: antisense morpholino oligo;
p190: p190RhoGAP-A;
PI(4,5)P_2_: phosphatidylinositol 4,5-bisphosphate;
ROI: region of interest;
SEM: Standard Error of the Mean;
ZnUMBA: zinc-based ultrasensitive microscopic barrier assay

## Acknowledgments

We thank all current and former members of the Miller lab for providing helpful discussions and feedback on this project. Specifically, we acknowledge Rachel Stephenson and Sara Varadarajan for their helpful insight and guidance on image acquisition and Rho flare quantification techniques. We also thank William Bement for pCS2+/mCherry-2xrGBD.

This work was funded by the National Institutes of Health (R01 GM112794 and R35 GM153204) to A.L. Miller, and a T32 fellowship from the University of Michigan Center for Cell Plasticity and Organ Design (T32 HD007505) to Z. Craig.

The authors declare no competing financial interests.

## Author contributions

Z. Craig, T.R. Arnold, and A.L. Miller conceptualized the study; Z. Craig and A.L. Miller developed the methodology; Z. Craig performed the majority of experiments and data analysis; T.R. Arnold performed experiments for Figure 1C and S2C; K. Walworth performed experiments for Figure 3B; A. Walkon performed the cloning and optimized concentration for the mNeon-p190RhoGAP-A construct for Supplemental Figure 2A and B; Z. Craig and A.L. Miller wrote the original draft of the manuscript; Z. Craig, A.L. Miller, and the other authors revised the manuscript; Z. Craig and A.L. Miller acquired funding; A.L. Miller supervised the study.

## Supplemental Figure Legends

**Figure S1. Anillin colocalizes with active RhoA at Rho flares and regulates junctional RhoA activity**

(A) Schematic of quantification approach for measuring active Rho, ZO-1, or Anillin intensity at Rho flares. A circular ROI (dashed green line) is placed on the junction undergoing a Rho flare (green shading) to measure the intensity of active Rho/ZO-1/Anillin. The same sized ROI is then placed on a reference junction to measure the intensity of active Rho/ZO-1/Anillin at a control region. The intensities at the Rho flare are divided by the intensities at the reference junction to normalize intensity.

(B) The Anillin morpholino targets the 5’ UTR of Anillin mRNA.

(C) Microscopy images showing whole-field view of ZO-1 intensity (Fire LUT) and Active RhoA intensity (Gray LUT) in control versus Anillin KD embryos. Images were acquired using the same microscopy settings, and LUT levels were adjusted the same.

(D) Overall intensity of active Rho at Rho flares (measured by the area under the curve) is not significantly different (p = 0.717) in control vs. Anillin KD Rho flares. Scatterplot showing area under the curve (calculated from **Figure 2C**). Error bars represent mean +/- SEM. n = 4 experiments, 7 embryos, 16 control flares, 14 Anillin KD flares.

Related to Figures 1 and 2.

**Figure S2. Anillin KD reduces actomyosin intensity**

(A) Microscopy images showing whole-field view of ZO-1 intensity (Gray LUT), Active RhoA intensity (Gray LUT), and F-actin intensity (Fire LUT) in control versus Anillin KD embryos. Images were acquired using the same microscopy settings, and LUT levels were adjusted the same.

(B) Averaged F-actin intensity line scans (control, light blue line vs. Anillin KD, dark blue line), shading represents SEM (calculated from plots like the representative plot in **Figure 3C**). (B’) Overall intensity of F-actin at Rho flares (measured by the area under the curve from data in **Figure S1B**) is not significantly different (p = 0.145) in control vs. Anillin KD embryos. Error bars represent mean +/- SEM. n = control: 1 experiment, 1 embryo, 5 control flares; Anillin KD: 2 experiments, 2 embryos, 10 Anillin KD flares.

(C) Microscopy images showing whole-field view of ZO-1 intensity (Gray LUT), Active RhoA intensity (Gray LUT), and Myosin II intensity (Fire LUT) in control versus Anillin KD embryos. Images were acquired using the same microscopy settings, and LUT levels were adjusted the same.

Related to Figure 3

**Figure S3. Anillin KD increases multicellular barrier leaks**

(A) Temporal color-code projection showing active RhoA intensity over 27 minutes. See Materials and Methods for details. (B) Frequency of barrier leaks at bicellular junctions (Bicellular Leaks/Junction/Second x 10^-5^) appears somewhat increased in Anillin KD embryos vs. controls, but this difference is not statistically significant (p = 0.158). Error bars represent mean +/- SEM. N = 3 experiments, 4 control embryos, 7 Anillin KD embryos.

Related to Figure 4.

**Figure S4. Anillin and p190RhoGAP-A colocalize at Rho flares.**

(A) Microscopy image shows that Anillin (Anillin-Halo with JF 646) and p190RhoGAP-A (mNeon-p190RhoGAPA) colocalize with active Rho (mCherry-2xrGBD) at the peak of Rho intensity during a Rho flare (magenta arrowheads). Yellow dashed box in ZO-1 channel (BFP-ZO-1) shows zoomed region for montage in (B).

(B) Time-lapse montage of Anillin, p190RhoGAP-A, active RhoA probe, and BFP-ZO-1, all shown in Fire LUT. Time=0 indicates the start of the Rho flare. Reduction in ZO-1 signal (open blue arrowhead) precedes Rho flare (yellow arrowhead, active Rho channel), which is accompanied by an increase in Anillin and p190RhoGAP-A (yellow arrowheads, Anillin and p190RhoGAP-A channels). Following the Rho flare, ZO-1 intensity is reinforced (solid blue arrowhead).

(C) Anillin intensity (Anillin-3xmCherry, grayscale) increases and decreases with similar timing to p190RhoGAP-A intensity (3xGFP-p190Rho-A, FIRE LUT) over the lifetime of a Rho flare. Related to Figure 1.

### Video Legends

**Video S1:** Time-lapse confocal imaging of gastrula-stage *Xenopus laevis* embryo shows that ZO-1 (BFP-ZO-1), active RhoA (mCherry-2xrGBD) and Anillin (Anillin-Halo with JF 647), all shown in Fire LUT, colocalize during a Rho flare. Video: time interval = 15 s, frame rate = 7 fps. Related to Figure 1, B and C.

**Video S2:** Time-lapse confocal imaging shows Rho flares in control (left) vs. Anillin KD (right) *Xenopus laevis* embryos. Anillin KD embryo has decreased overall junctional active RhoA intensity (GFP-rGBD, grayscale), and Rho flare exhibits (white arrow) increased intensity over background and shorter duration compared with control Rho flare (white arrow). Anillin KD embryos fail to reinforce their tight junctions (BFP-ZO1, FIRE LUT) following the Rho flare. Video: time interval = 15 s, frame rate = 12 fps. Brightness and contrast were adjusted to the same levels for both videos. Related to Figure 2, A-D’.

**Video S3:** Time-lapse confocal imaging shows F-actin at Rho flares in control (left) vs. Anillin KD (right) *Xenopus laevis* embryos. Compared with controls, which have broad F-actin accumulation (LifeAct-mRFP, FIRE LUT) associated with the Rho flare (GFP-rGBD, grayscale), Anillin KD embryos have intense F-actin signal that is confined at the junction at Rho flare sites. Video: time interval = 15 s, frame rate = 12 fps. Gray LUT adjusted to 0-1451 for controls and 77-2699 for Anillin KD. Fire LUT adjusted to 95-1569 for controls and 59-4095 for Anillin KD. Related to Figure 3, A and C.

**Video S4:** Time-lapse confocal imaging showing Myosin II at Rho flares in control (left) vs. Anillin KD (right) *Xenopus laevis* embryos. Compared with controls, Anillin KD embryos have reduced intensity of Myosin II signal overall (SF9-mCherry, FIRE LUT) and fail to accumulate Myosin II at Rho flares (GFP-rGBD, grayscale). Video: time interval = 15 s, frame rate = 12 fps. Related to Gray LUT adjusted to 186-1350 for controls and 77-1714 for Anillin KD. Fire LUT adjusted to 10-892 for controls and 77-1313 for Anillin KD. Figure 3, B and D-F’.

**Video S5:** Time-lapse confocal imaging showing barrier function leaks visualized by changes in FluoZin3 intensity (Fire LUT) in control (left) vs. Anillin KD (right) embryos. Anillin KD embryos have increased baseline FluoZin3 intensity and increased frequency of tight junction leaks. Video: time interval = 10 s, frame rate = 15 fps. Brightness and contrast were adjusted to the same levels for both videos. Related to Figure 4, A-F.

