## Supplementary figures and images for "Anillin tunes contractility and regulates barrier function during Rho flare-mediated tight junction remodeling"

### Supplemental Figures

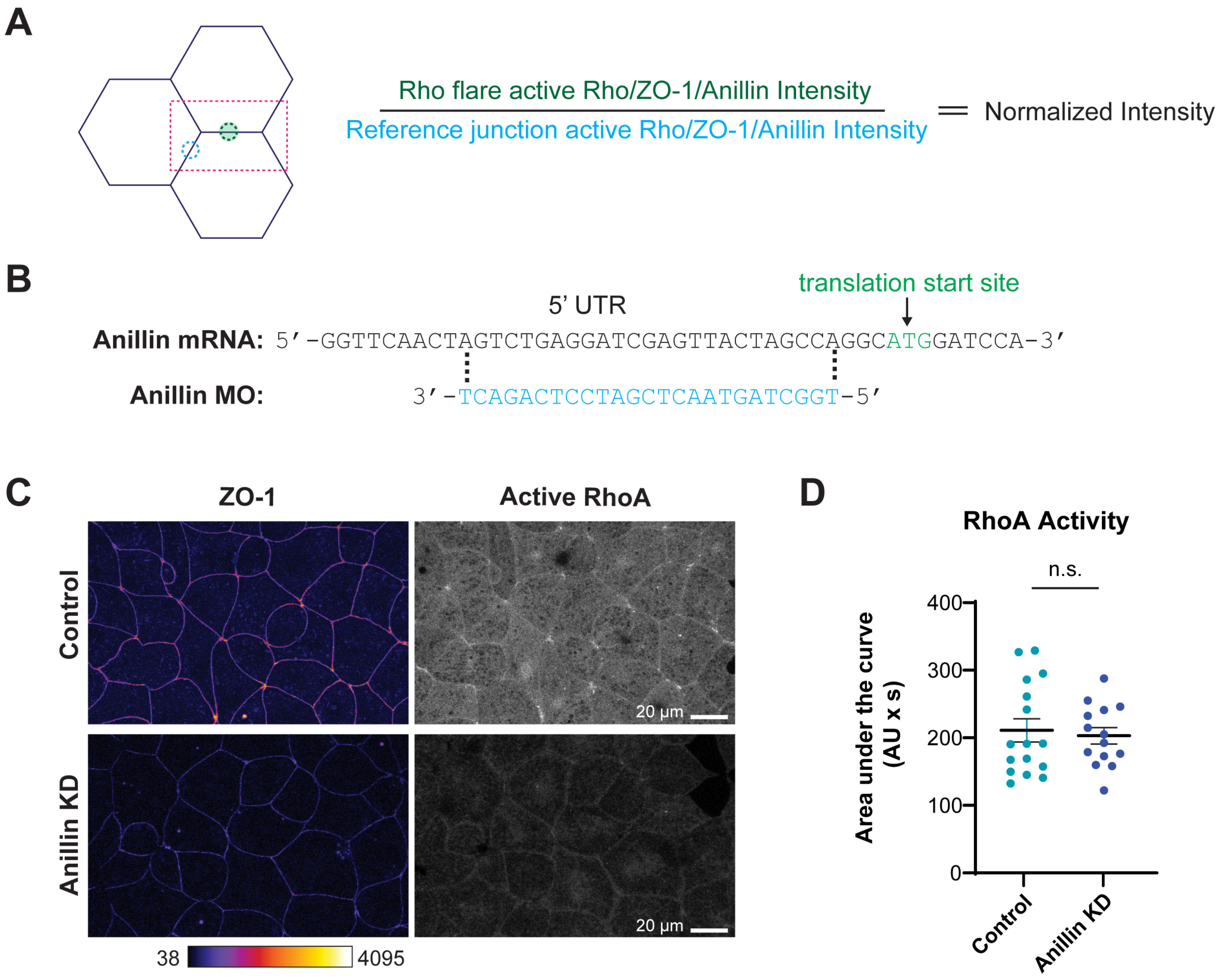

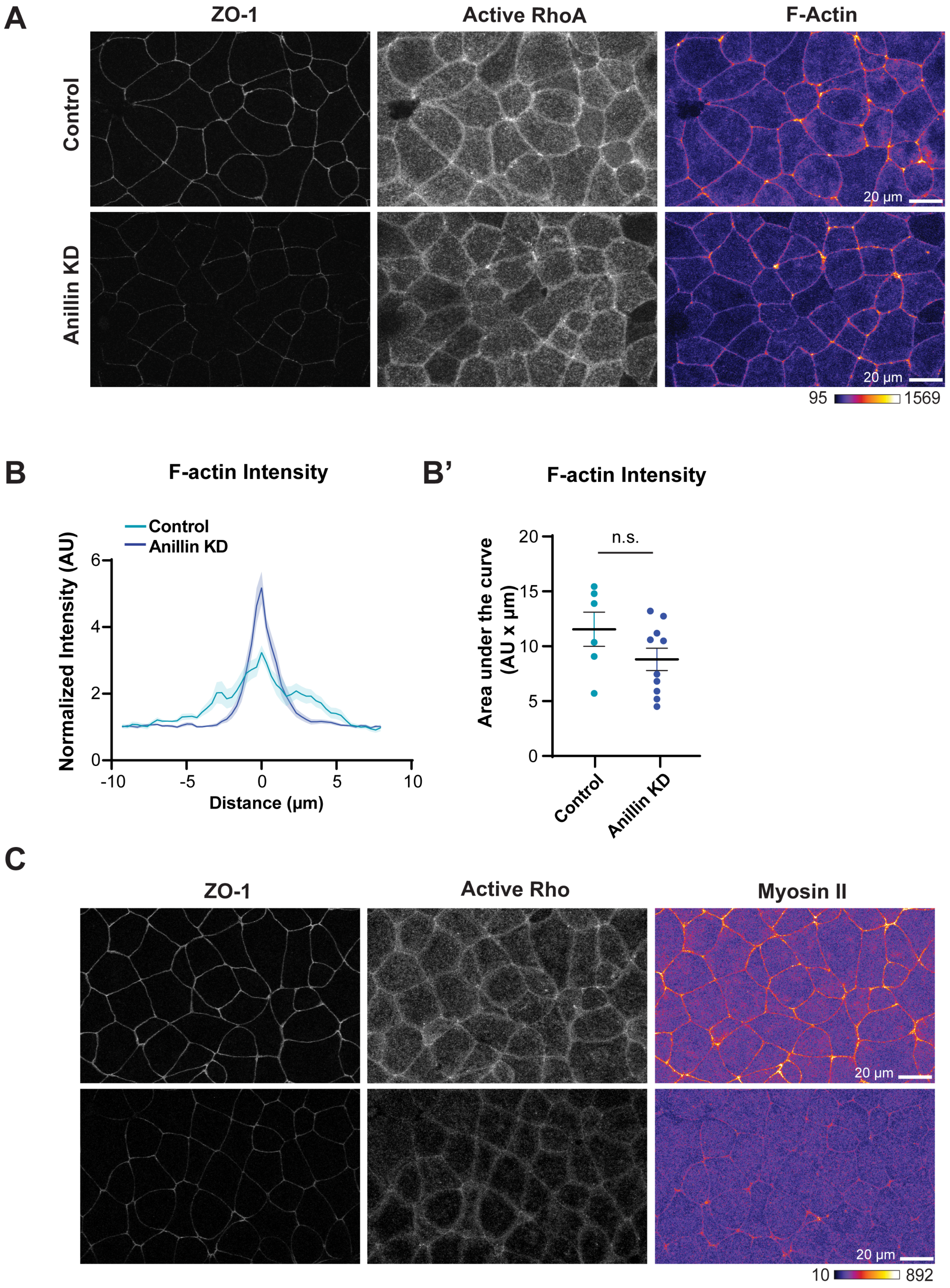

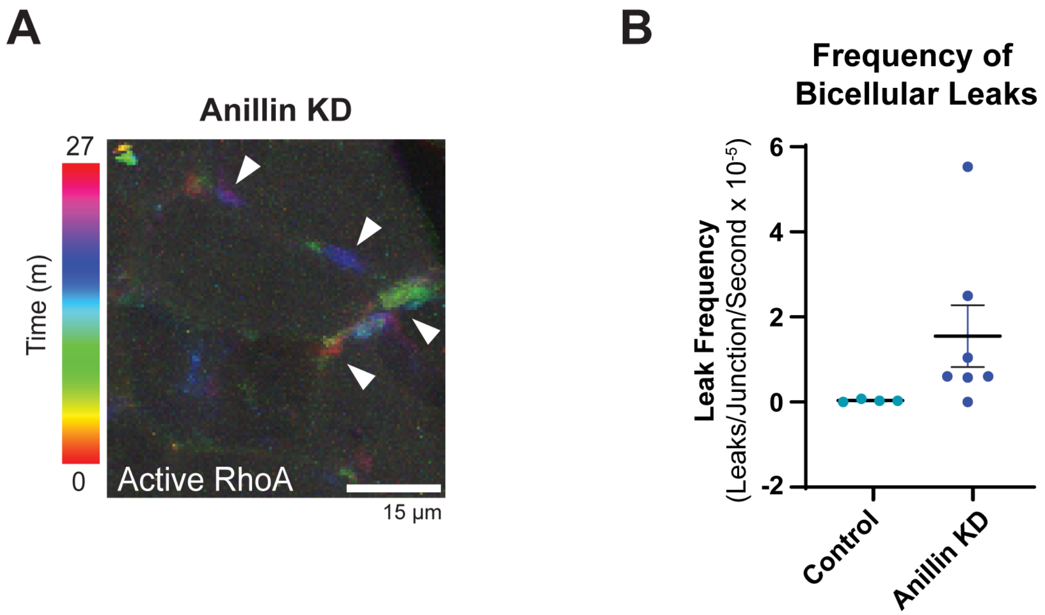

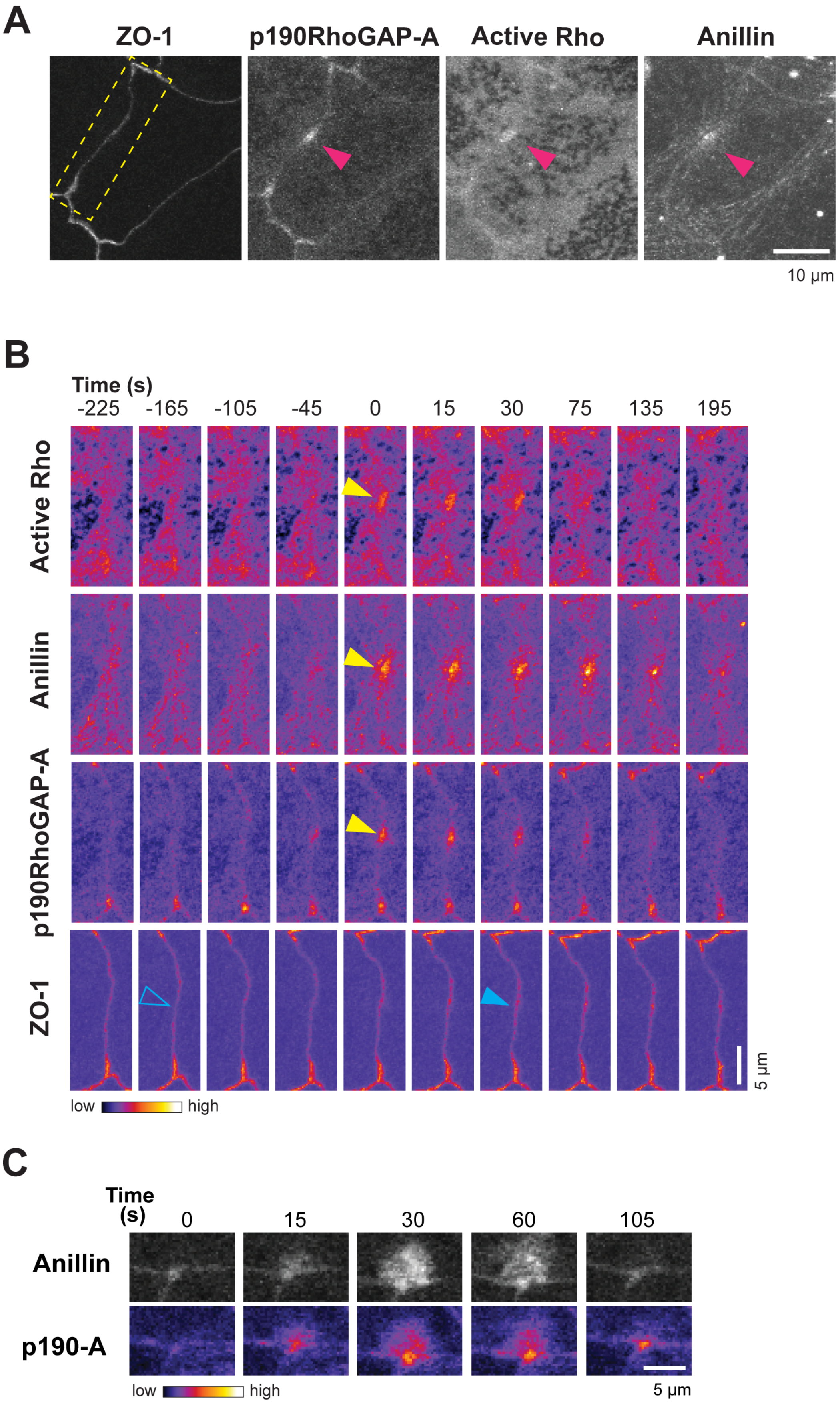
